# Composition and short chain fatty acid formation of pig fecal microbiota during growth on predigested wheat and rye *in vitro*

**DOI:** 10.1101/2025.07.12.664509

**Authors:** Clara Berenike Hartung, Sabrina Woltemate, Andreas von Felde, Knud Erik Bach Knudsen, Christian Visscher, Marius Vital

## Abstract

**Background:** While wheat is the most commonly used cereal in pig feeding, rye has gained renewed interest as a component in pig diets due to its positive effects on gut health. The effects of distinct cereal sources on the composition and functionality of gut microbiota are, however, poorly investigated so far. *In vitro* studies offer a valuable approach to gain detailed insights on this matter, however, require a substrate that closely mimics the conditions found in the large intestine. In this study, we performed predigestion of cereal substrates using ileo-cecally fistulated minipigs simulating the upper gastrointestinal digestion. The composition of microbiota, total cell counts, and short chain fatty acid concentrations after a 24-hour incubation were analysed. Results were compared to cultures containing native substrates.

**Results:** Both acetate and propionate concentration were significantly higher in predigested rye compared to predigested wheat (8.17 ± 2.71 vs. 6.55 ± 2.16 mmol/l and 3.40 ± 0.436 vs. 2.64 ± 0.337 mmol/l, respectively), whereas butyrate concentrations were not different. Substantial differences were observed between native and predigested substrates with a higher butyrate production on the former. Bacterial cell counts were higher in predigested substrates and elevated in rye cultures compared to respective wheat incubations. Bacterial communities differed markedly between the two predigested cereals as well as between native and predigested substrates. Several known butyrate and propionate producers – for example *Agathobacter, Blautia, Coprococcus, Mediterraneibacter* and *Roseburia* – grew significantly more on predigested rye than on predigested wheat. *Bifidobacterium, Clostridium sensu stricto,* and *Succinivibrio* were significantly more abundant on native substrates compared to their predigested forms.

**Conclusion:** Rye promoted higher production of acetate and propionate compared with wheat along with a distinct bacterial community composition. Predigestion of substrates was crucial in *in vitro* experiments for gaining adequate insights into growth and substrate degradation by colonic microbiota.

## Introduction

Wheat is the standard substrate for studies investigating the impact of different cereal grains on the large intestinal environment of pigs, given its widespread use as the primary grain in pig diets throughout Europe [1]. However, rye has recently come into focus of swine feeding [2–7] as it is a durable crop [8] and healthy feed source [9, 10]. Rye has been proven to alter the microbial community in pigs in a *Salmonella* infection challenge with promoted growth of *Bifidobacterium*, several lactic acid bacteria and *Faecalibacterium prausnitzii* in the cecum [11] along with reduced *Salmonella* shedding [10]. Rye has the highest content of soluble non-starch polysaccharides (NSP) among the common cereal sources [12–14]. Due to their chemical structure, such as molecular bonds between carbohydrate sugar residues, a large part of the NSP escape digestion in the small intestine [15] characterizing them as dietary fiber. Dietary fiber is defined as carbohydrate polymers with a degree of polymerization of at least three, which are neither digested nor absorbed in the small intestine and therefore reach the large intestine [15, 16] where they can be used by microbiota as substrates [17] impacting host physiology. A large proportion of the NSP in rye consists of fructans and arabinoxylan [18, 19], which are primarily converted into the short chain fatty acids (SCFA) acetate, propionate and butyrate [20]. Propionate has positive effects on the lipid metabolism [21] and the secretion of satiety related hormones [22] of pigs. Butyrate stabilizes the intestinal barrier function [23, 24] promoting gut health and enhances overall growth performance.

Rye and wheat differ not only in their content of soluble NSP but also in total NSP, starch, protein, amino acids and overall energy content [12]. Additional differences are seen in the ileal digestibility in pigs [25], hence, increasing the differences in composition of substrates when reaching the large intestine. The carbohydrate fraction of rye displays a lower prececal digestibility than that of wheat bringing more fermentable substrate to colonic microbiota [9]. Proteins of rye show lower prececal digestibility in pigs compared to wheat as well [25], causing higher amounts of protein entering the hindgut. Fermentation of non-digested protein in the large intestine is often found to lead to the formation of potentially harmful nitrogen containing bacterial metabolites [26], however, can also be used by some bacteria for the formation of propionate and butyrate [27].

*In vitro* experiments are used to simulate physiological processes that are difficult to investigate *in vivo*. For instance, to study the environment in the large intestine and the processes involving microbiota *in vivo*, either feces from live animals or digesta from slaughtered animals has to be analysed. However, obtained material only provides snapshots and cannot depict the dynamic processes involved. For example, true production of SCFA in the gut is not reflected by SCFA concentration in intestinal material as SCFA are rapidly absorbed and metabolized by the host [28] uncoupling SCFA concentrations measured in feces and the circulation from actual SCFA production. *In vitro* experiments, however, allow to investigate substrate degradation, metabolite formation and associated changes in microbiota in detailed quantitative terms [29]. In this context, for closely mimicking *in vivo* conditions, *in vitro* experiments need to accurately mirror conditions in the large intestine [30]. A significant factor in this context is substrate composition [31, 32] and to determine the influence of feed on colonic microbiota, substrate composition should be as close as possible to conditions the bacteria experience *in vivo*. For example, cereal grains contain high levels of starch [12], which is almost completely digested in the small intestine [33] and only a limited extent reaches the colon. For *in vitro* research, it is therefore recommended that the grain is pre-treated chemically with a mixture of hydrochloric acid, pepsin, and small intestinal enzymes [34, 35]. In this study we used an alternative approach, namely, the effluent from ileal fistulated pigs that can supply an *in vivo* “predigestion”, which provides substrate with a composition similar to what can be experienced *in vivo* as next to substrate degradation its digested parts are also actively absorbed by the host. We investigated the influence of predigested wheat and predigested rye on the intestinal microbiota of pigs and their ability to synthesize SCFA *in vitro.* Results were compared to those derived from native substrate incubations in order to examine the impact of substrate predigestion on bacterial growth. We hypothesised that predigested rye would lead to a higher production of SCFA and an alteration in microbial community composition and that predigestion of substrates lead to different outcomes compared to incubations with their native forms

## Materials and Methods

### In vitro incubations

For *in vitro* incubations, 12 healthy pigs (conventional fattening hybrids, six female and castrated male each, 22.8 ± 10.0 kg bodyweight) were available. Pigs were housed in pairs in pens of 3 x 1 m and were fed twice daily with complete diets to provide twice the energy for maintenance requirements. Diets contained either 69 % of wheat or rye, respectively, mixed with extracted soybean meal, barley, potato protein, calcium carbonate, monocalcium phosphate, soybean oil, sodium chloride and a vitamin-/mineral-mix in both diets (Raiffeisen Kraftfutterwerk Mittelweser Heide GmbH, Schweringen, Germany) to adapt the pigs to each cereal source before the *in vitro* experiments. Each pair of pigs received both diets consecutively for 11 days respectively with a “wash out” phase of seven days in between, during which a commercial piglet diet was fed (Deutsche Tiernahrung Cremer GmbH & Co. KG, Düsseldorf, Germany). On the eleventh day of the respective feeding phase, feces were collected directly after defecating and immediately transferred to a vinyl anaerobic chamber (Coy Laboratory Products, Grass Lake, MI, USA). Two-hundred mg of feces were diluted with 19.8 ml of 1x PBS and filtered through a 30 µm filter (Miltenyi Biotec, Bergisch Gladbach, Germany). In Hungate tubes (sealed with butyl rubber stoppers and screw caps (Chemglas Life Sciences, Vineland, New Jersey, USA)), 10 ml of an anaerobic medium (modified according to Reichardt et al. [36] and Kircher et al. [37] without bile acids and only acetate as SCFA), 20 mg of the respective substrate (2 g l^−1^ final concentration) and 100 µl of filtered feces suspension (∼5 × 10^7^ bacterial ml^−1^) were put together. Depending on the feeding of the pig, four tubes were fitted with each substrate. Substrates were native ground rye (if the pig was fed the diet containing 69 % rye) and wheat (if the pig was fed the diet containing 69 % of wheat) as well as predigested rye and predigested wheat (independent of the feeding of the pig); resulting in 12 tubes per pig. Two tubes per substrate were incubated for 24 hours at 37 °C (200 rpm; Edmund Bühler GmbH, Bodelshausen, Germany). From the other two tubes, samples were directly taken to receive values pre-incubation (“zero values”). Samples were taken for analysis of SCFA concentration, flow cytometry and 16S rRNA gene analysis (see below).

### Collection of predigested substrate for the in vitro incubation

Three adult Göttingen minipigs (Ellegaard®, two female, one castrated male, 31.8 ± 2.40 kg bodyweight) were used to generate substrates for *in vitro* incubations. All animals were fitted with an ileo-cecal re-entrant fistula which had been surgically implanted according to a modified method of Easter and Tanksley [38]. The fistula allowed the complete collection of ileal digesta. The pigs were fed 100 % of ground wheat or rye (each pig received both cereals with a break of at least seven days in between) in portions of 250 g (+ 1 l tab water) twice a day for two days. On the third day, after the morning meal, the fistulas were opened and the total mass of ileal digesta was collected for nine hours. The collected material was then pooled, lyophilised, ground and stored at 4° C. Before using the material in *in vitro* incubations, 100 % ethanol was added for at least 10 minutes to eradicate the residual ileal microflora and then removed by drying in a vacuum dryer.

### Analysis of the native and predigested substrate material

Nutrient contents (starch, crude protein, amino acids) of the ground cereals and predigested material were analysed according to standard methods by VDLUFA [39] in duplicate. The dry matter (DM) content was measured by drying the samples at 103 °C until constant weight was reached. The starch content was determined by polarimetric measurement (Polatronic E®, Schmidt + Haensch GmbH & Co., Berlin, Germany). The total nitrogen (N) content was examined using the DUMAS combustion method (rapid MAX N exceed, Elementar, Analysen-Systeme GmbH, Hanau, Germany). Total nitrogen was multiplied by a constant factor of 6.25 to calculate the crude protein (CP) content. The content of amino acids was determined via ion exchange chromatography (Biochrom 30+, Biochrom GmbH, Berlin, Germany). The non-starch polysaccharides (NSP) and dietary fibre (DF) were analysed via gas chromatography according to a method described by Bach Knudsen et al. 1997 [40].

### SCFA Analysis

For SCFA analysis, before and after 24 hours of incubation, 100 µl of the suspension was added to 900 µl of purified water and 100 µl of an internal standard (10 ml 17 % phosphoric acid, 0.025 ml 4-methyl valeric acid). Separation of the SCFA components was done on a 30 m column (Supelco, NukolTM, Capillary GC Column) in a gas chromatograph (GC-2014, Shimadzu, Duisburg). The column temperature was 180 °C (injector temperature 220 °C, detector temperature 250 °C). The SCFA were determined using a flame ionization detector during a 25-minute analysis time in the following order: acetic acid, propionic acid, i-butyric acid, n-butyric acid, i-valeric acid, n-valeric acid with reference to the internal standard. The detection limit value for individual SCFA was 0.01 mmol kg^−1^. The production of SCFA *in vitro* was calculated by subtracting the “zero value” from the value analysed after 24 hours of incubation.

### Flow Cytometry

For flow-cytometric measurements (FCM), 1 ml material from the Hungate tubes was taken before and after 24 hours of incubation and filtered through a 30 µm filter (Miltenyi Biotec, Bergisch Gladbach, Germany). The filtrate was then diluted with 1x PBS (fivefold for samples before incubation, 500-fold for samples after incubation). All suspensions were stained with 10 µl EDTA and 10 µl SYBR Green (Thermo Fisher Scientific, United States) according to Hammes et al. [41] and incubated for 15 min in the dark. Stained samples were diluted 10-fold in 1x PBS before measurements and cell concentrations were recorded on a MACSQuant Analyzer 10 (Miltenyi Biotec, Germany). The growth of bacterial cells was calculated by subtracting the “zero value” from the values found after 24 hours of incubation.

### 16S rRNA gene analysis

Before and after incubations, 2 ml of the suspension were centrifuged (15.400 g, 4 °C) and the pellet was used for DNA extraction. DNA was extracted (ZymoBIOMICS DNA Miniprep Kit, Zymo Research Europe GmbH, Freiburg, Germany) including a beat-beating step (5x 1 min with 5 min rest between steps on FastPrep-24 5G at speed 5.5). Amplification of the V3V4 region of the 16S rRNA gene and subsequent sequencing on Illumina MiSeq (2x 250 bp; San Diego, CA, USA) was done as before [42].

### Statistical Analysis

Data were compared for differences between incubations with the predigested rye and predigested wheat and between the incubations of the respective cereals in native and predigested forms.

Data for SCFA and cell counts were analyzed using SAS v. 7.1 (SAS Inst. Inc. Cary., NC, USA) and examined for their normal distribution using the Kolmogorov-Smirnov test and Shapiro-Wilk test. For group comparison for normally distributed data, the T-test was applied and for non-normal data the unpaired two-samples Wilcoxon test. The significance level was specified with p < 0.05.

For 16S rRNA gene data analyses were performed in R (v4.2.2) applying linear models (function *lm*) using log transformations (log(data+1)). All taxa with an average abundance of 0.1% were included; for absolute abundance data proportional count data were multiplied by bacterial concentrations. All results were corrected for multiple testing using *fdrtools* (v1.2.17), where a lfdr <0.05 was considered significant. Ordination analyses are based on metric multidimensional scaling (MDS) using the *phyloseq* package (v1.36.0).

## Results

The feeding of the donor pigs before the *in vitro* experiment had no significant effects on any parameter measured (SCFA concentration, cell growth and community composition) and results were, therefore, only compared between individual substrates.

### Composition of the native and predigested substrates

Analysis of the substrates were performed in duplicates in pooled samples. The crude protein content in wheat relatively decreased during predigestion, whereas it increased in rye (Table 1) with values being generally lower in rye. Out of 17 analysed amino acids, 11 relatively increased in predigested rye, whereas only six relatively increased in predigested wheat compared to the native substrate.

**Table 1.** Nutrient content of the native and predigested cereals (% of DM).

| % DM | Wheat | Predigested Wheat | Relative change from native to predigested | Rye | Predigested Rye | Relative change from native to predigested |
| --- | --- | --- | --- | --- | --- | --- |
| Crude protein | 14.3 | 12.4 | ↓ | 8.12 | 10.4 | ↑ |
| Aspartate | 0.713 | 0.980 | ↑ | 0.626 | 0.824 | ↑ |
| Threonine | 0.408 | 0.544 | ↑ | 0.303 | 0.468 | ↑ |
| Serine | 0.646 | 0.593 | ↓ | 0.370 | 0.489 | ↑ |
| Glutamine | 4.57 | 1.27 | ↓ | 2.03 | 1.04 | ↓ |
| Glycine | 0.566 | 1.16 | ↑ | 0.383 | 1.05 | ↑ |
| Alanine | 0.497 | 0.747 | ↑ | 0.369 | 0.534 | ↑ |
| Valine | 0.607 | 0.610 | ↑ | 0.422 | 0.505 | ↑ |
| Cysteine | 0.304 | 0.236 | ↓ | 0.130 | 0.182 | ↑ |
| Methionine | 0.219 | 0.153 | ↓ | 0.135 | 0.116 | ↓ |
| Isoleucine | 0.490 | 0.409 | ↓ | 0.280 | 0.348 | ↑ |
| Leucine | 0.917 | 0.696 | ↓ | 0.523 | 0.533 | ↑ |
| Tyrosine | 0.401 | 0.301 | ↓ | 0.206 | 0.243 | ↑ |
| Phenylalanine | 0.621 | 0.418 | ↓ | 0.365 | 0.337 | ↓ |
| Histidine | 0.318 | 0.231 | ↓ | 0.194 | 0.194 | - |
| Lysine | 0.388 | 0.428 | ↑ | 0.321 | 0.393 | ↑ |
| Arginine | 0.654 | 0.524 | ↓ | 0.445 | 0.425 | ↓ |
| Proline | 1.40 | 0.579 | ↓ | 0.697 | 0.542 | ↓ |

Both cereals lost most of the starch content during small intestinal digestion. The content of fructan, soluble NSP, DF, arabinose and xylose was higher in native rye than native wheat (see Table 2). That difference became even more pronounced in predigested cereals, where the relative contents for those compounds increased notably. Relative contents of cellulose and lignin were higher in native wheat and predigested wheat than in respective rye substrates.

**Table 2.**
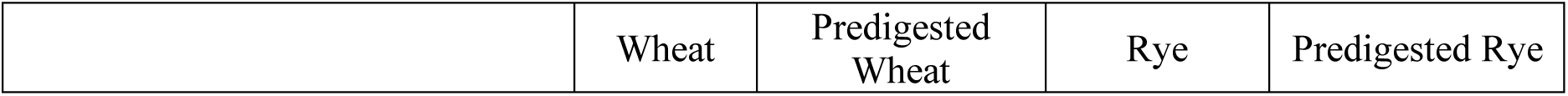
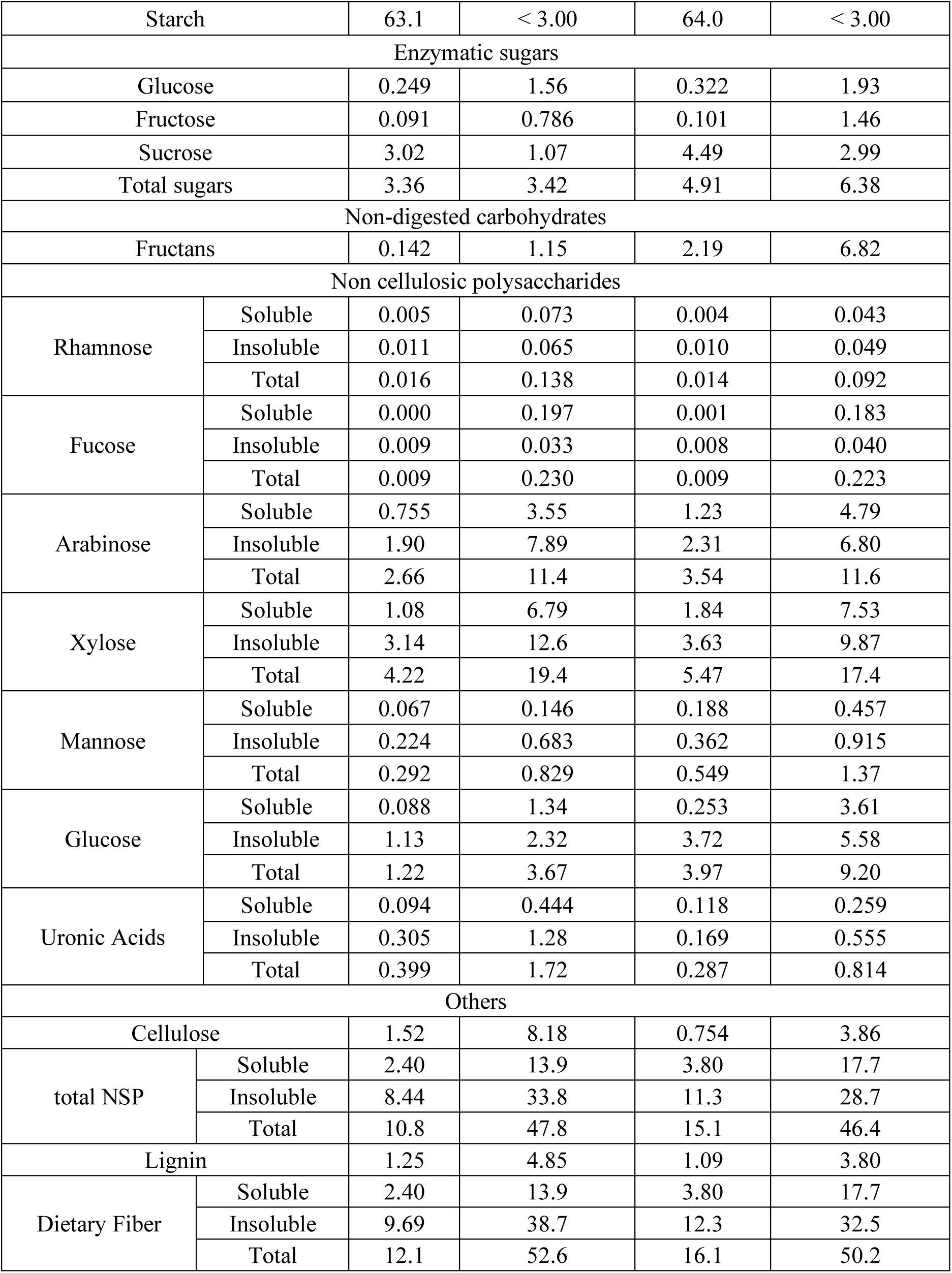
Carbohydrate and lignin content of the native and predigested cereals (% of DM)

|  | Wheat | Predigested Wheat | Rye | Predigested Rye |
| --- | --- | --- | --- | --- |

|  |  |  |  |  |  |
| --- | --- | --- | --- | --- | --- |
| Starch |  | 63.1 | < 3.00 | 64.0 | < 3.00 |
| Enzymatic sugars |  |  |  |  |  |
| Glucose |  | 0.249 | 1.56 | 0.322 | 1.93 |
| Fructose |  | 0.091 | 0.786 | 0.101 | 1.46 |
| Sucrose |  | 3.02 | 1.07 | 4.49 | 2.99 |
| Total sugars |  | 3.36 | 3.42 | 4.91 | 6.38 |
| Non-digested carbohydrates |  |  |  |  |  |
| Fructans |  | 0.142 | 1.15 | 2.19 | 6.82 |
| Non cellulosic polysaccharides |  |  |  |  |  |
| Rhamnose | Soluble | 0.005 | 0.073 | 0.004 | 0.043 |
|  | Insoluble | 0.011 | 0.065 | 0.010 | 0.049 |
|  | Total | 0.016 | 0.138 | 0.014 | 0.092 |
| Fucose | Soluble | 0.000 | 0.197 | 0.001 | 0.183 |
|  | Insoluble | 0.009 | 0.033 | 0.008 | 0.040 |
|  | Total | 0.009 | 0.230 | 0.009 | 0.223 |
| Arabinose | Soluble | 0.755 | 3.55 | 1.23 | 4.79 |
|  | Insoluble | 1.90 | 7.89 | 2.31 | 6.80 |
|  | Total | 2.66 | 11.4 | 3.54 | 11.6 |
| Xylose | Soluble | 1.08 | 6.79 | 1.84 | 7.53 |
|  | Insoluble | 3.14 | 12.6 | 3.63 | 9.87 |
|  | Total | 4.22 | 19.4 | 5.47 | 17.4 |
| Mannose | Soluble | 0.067 | 0.146 | 0.188 | 0.457 |
|  | Insoluble | 0.224 | 0.683 | 0.362 | 0.915 |
|  | Total | 0.292 | 0.829 | 0.549 | 1.37 |
| Glucose | Soluble | 0.088 | 1.34 | 0.253 | 3.61 |
|  | Insoluble | 1.13 | 2.32 | 3.72 | 5.58 |
|  | Total | 1.22 | 3.67 | 3.97 | 9.20 |
| Uronic Acids | Soluble | 0.094 | 0.444 | 0.118 | 0.259 |
|  | Insoluble | 0.305 | 1.28 | 0.169 | 0.555 |
|  | Total | 0.399 | 1.72 | 0.287 | 0.814 |
| Others |  |  |  |  |  |
| Cellulose |  | 1.52 | 8.18 | 0.754 | 3.86 |
| total NSP | Soluble | 2.40 | 13.9 | 3.80 | 17.7 |
|  | Insoluble | 8.44 | 33.8 | 11.3 | 28.7 |
|  | Total | 10.8 | 47.8 | 15.1 | 46.4 |
| Lignin |  | 1.25 | 4.85 | 1.09 | 3.80 |
| Dietary Fiber | Soluble | 2.40 | 13.9 | 3.80 | 17.7 |
|  | Insoluble | 9.69 | 38.7 | 12.3 | 32.5 |
|  | Total | 12.1 | 52.6 | 16.1 | 50.2 |

### SCFA formation during in vitro incubations

Major differences were found between the different substrates (Figure 1). Comparing predigested rye and predigested wheat, the concentrations *in vitro* of acetate and propionate differed significantly with higher values for predigested rye (8.17 ± 2.71 vs. 6.55 ± 2.16 mmol l^−1^ and 3.40 ± 0.436 vs. 2.64 ± 0.337 mmol l^−1^, respectively). Butyrate concentrations were nearly the same for both substrates (2.06 ± 0.385 vs. 2.00 ± 0.296 mmol l^−1^ for predigested rye and predigested wheat, respectively). Comparisons between the respective cereal in the native and predigested form yielded significantly higher acetate values for predigested substrates, whereas butyrate was significantly higher with native substrates (rye: acetate 5.37 ± 1.76 vs. 8.17 ± 2.71 mmol l^−1^ and butyrate 3.17 ± 0.979 vs. 2.06 ± 0.385 mmol l^−1^; wheat: acetate 4.18 ± 2.15 vs. 6.55 ± 2.16 mmol l^−1^ and butyrate 3.23 ± 0.733 vs. 2.00 ± 0.296 mmol l^−1^). Propionate production differed significantly between native and predigested wheat (native 3.18 ± 0.492 mmol l^−1^ vs. predigested 2.64 ± 0.337 mmol l^−1^), but not between native and predigested rye.

**Figure 1.**
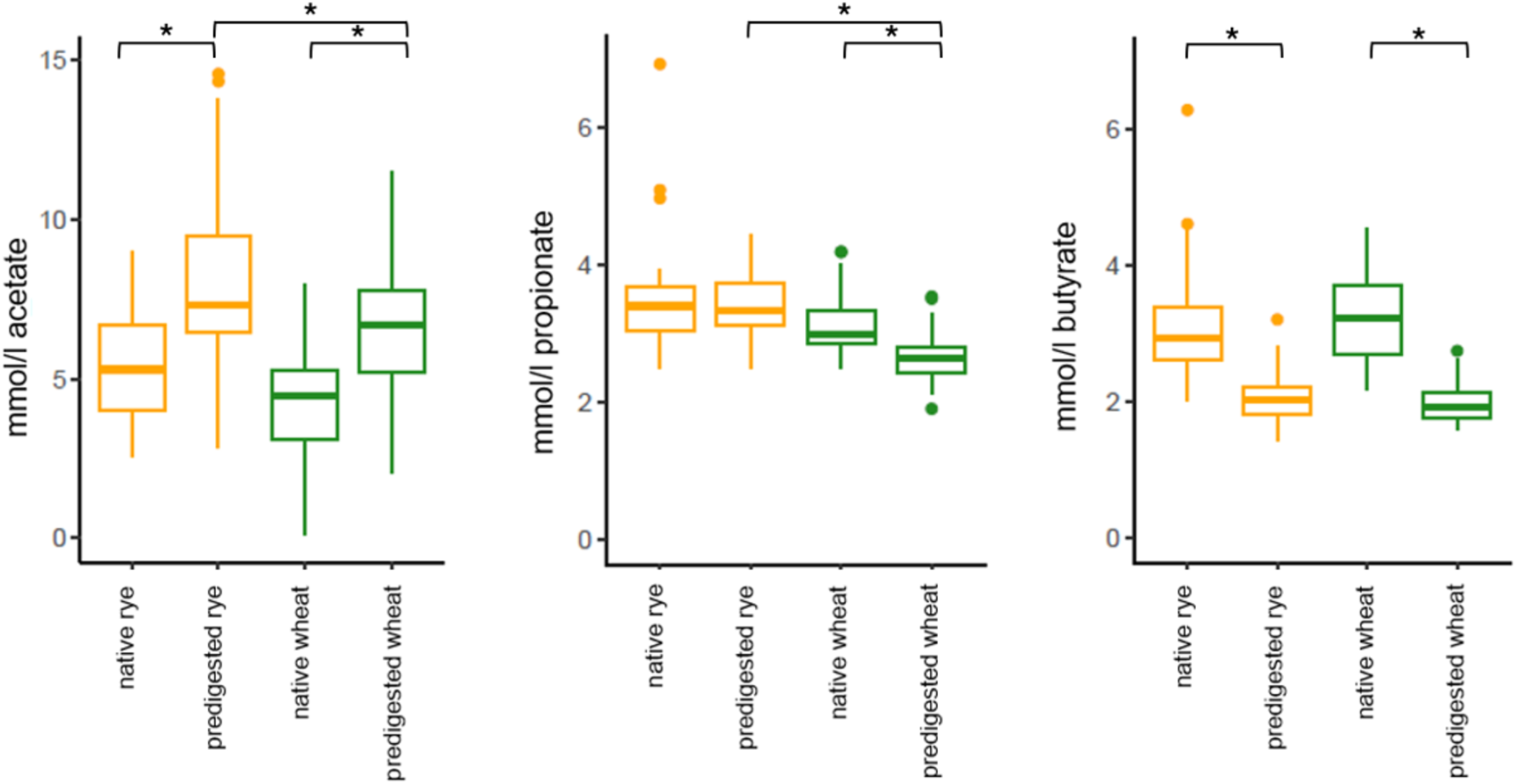
Acetate, propionate and butyrate (mmol/l) produced in 24 hours of *in vitro* incubation from different substrates incubated with pig fecal inocula (n=12); * indicate significant differences between respective substrates.

### Bacterial concentration (flow cytometry)

Significantly more cells were produced on predigested rye compared with predigested wheat (1.39 × 10^9^ ± 0.194 × 10^9^ ml^−1^ vs. 1.20 × 10^9^ ± 0.156 × 10^9^ ml^−1^), whereas there were no differences either between the two native substrates (native rye 1.34 × 10^9^ ± 0.37 × 10^9^ ml^−1^ vs. native wheat 1.16 × 10^9^ ± 0.18 × 10^9^ ml^−1^) nor between the respective native and predigested substrates (Figure 2).

**Figure 2.**
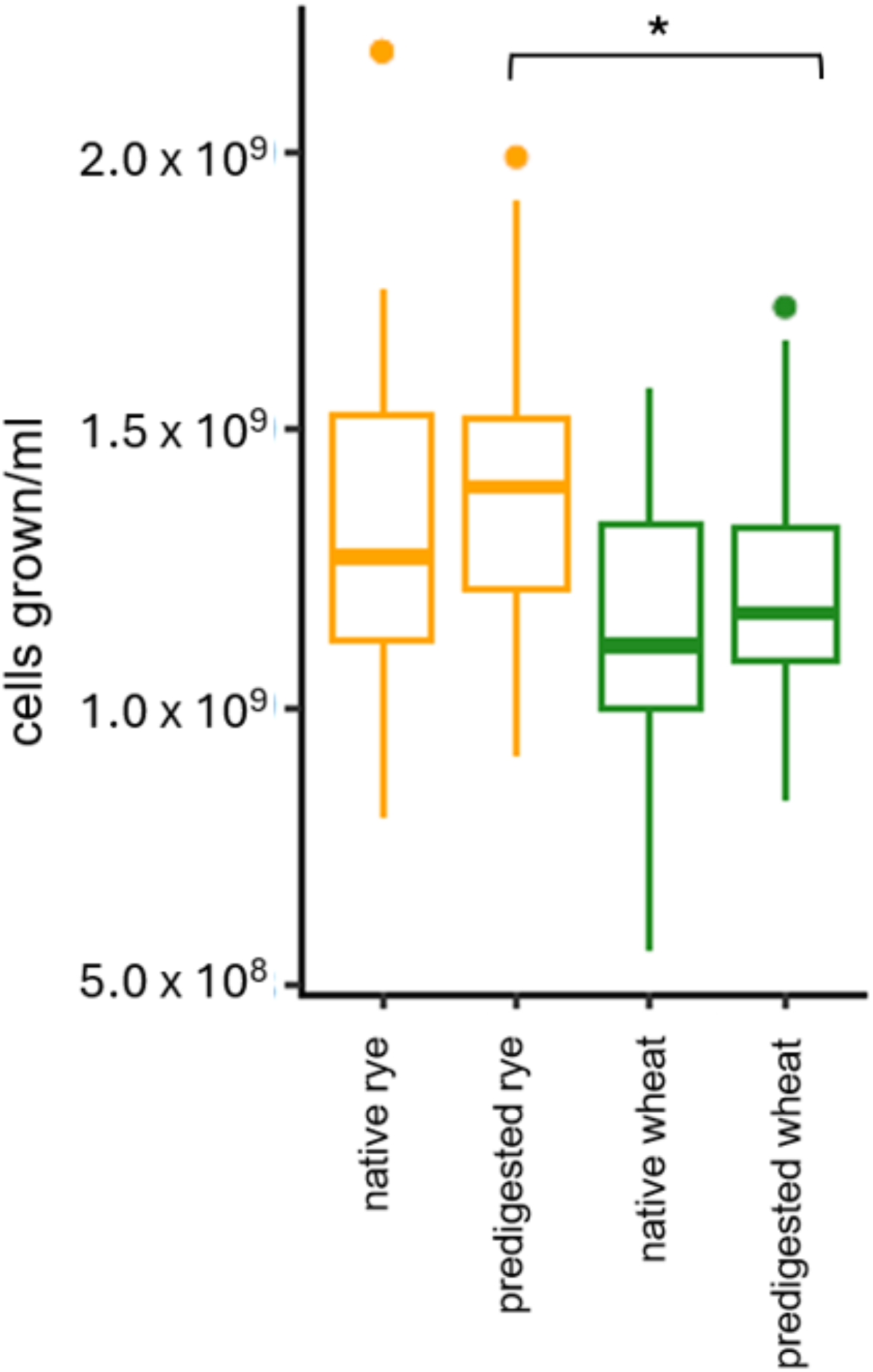
Number of cells (cells/ml) grown in 24 hours of *in vitro* incubation from different substrates incubated with pig fecal inocula (n=12); * indicate significant differences between respective substrates.

### Bacterial composition (16S rRNA gene analysis)

Ordination analysis of overall communities based on absolute bacterial numbers revealed a clear differentiation between communities grown on native and predigested substrates, respectively, but with no clustering between the respective cereals (Figure 3). Comparing only communities grown on predigested substrates a clear differentiation between rye and wheat was observed (Figure 4). Communities could clearly be matched to the corresponding predigested substrate, regardless of the donor animal. Individual differences between the donor pigs were also noted even though they were all housed and fed alike. For communities grown on native substrates, no clear differentiation between rye and wheat was found but clustering for each animal was observed (Figure 5).

**Figure 3.**
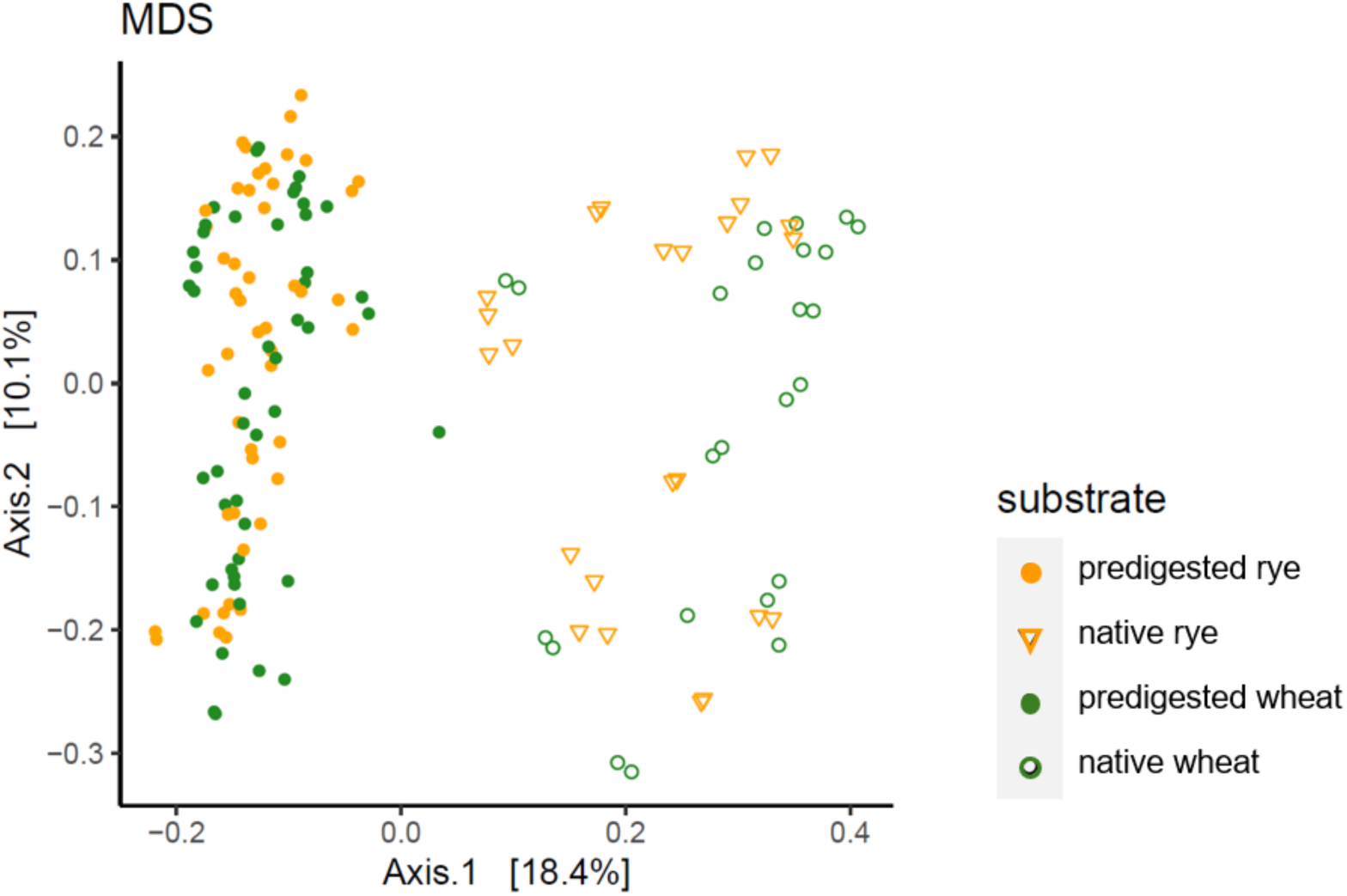
Ordination analysis of bacterial communities (absolute abundances) grown on all substrates (native and predigested rye and wheat) after 24 hours from pig fecal inocula (n=12).

**Figure 4.**
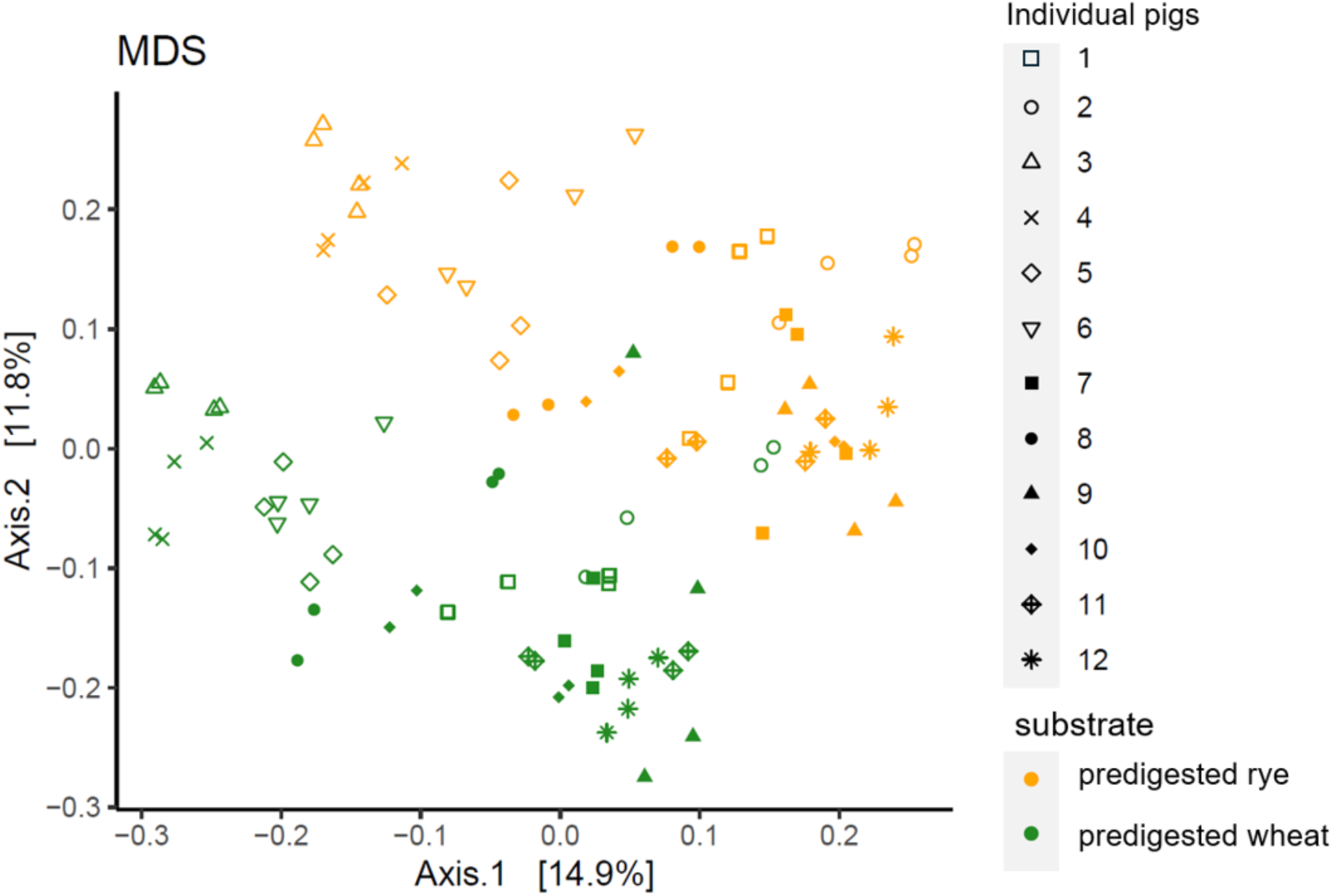
Ordination analysis of bacterial communities (absolute abundances) grown on predigested rye and predigested wheat after 24 hours from pig fecal inocula (n=12).

**Figure 5.**
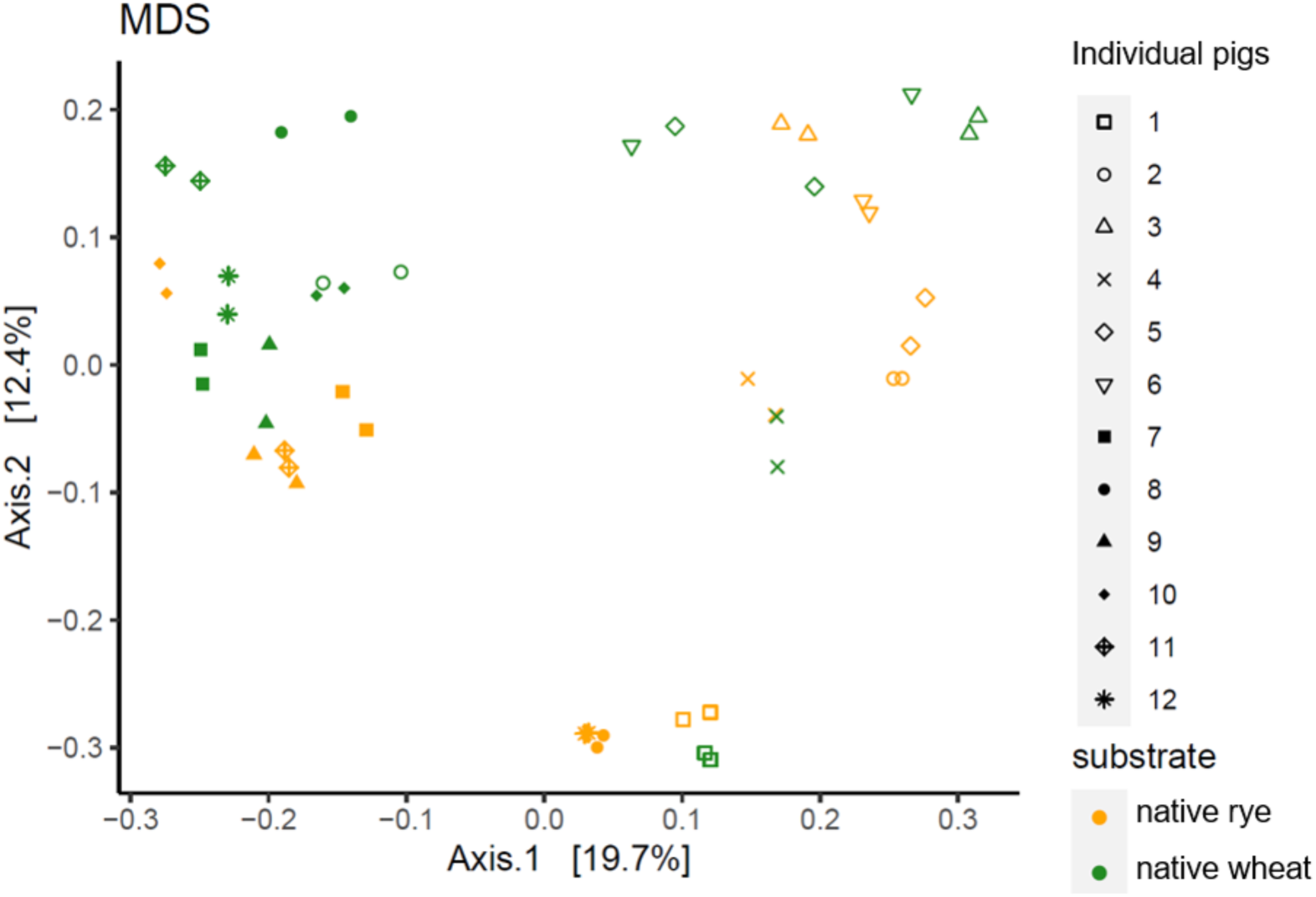
Ordination analysis of bacterial communities (absolute abundances) grown on native rye and native wheat after 24 hours from pig fecal inocula (n=12).

Significant differences were found in absolute abundances (Figure 6). On predigested rye, significantly more *Actinobacteria* and *Firmicutes* (1.36 × 10^8^ ml^−1^ vs. 0.863 × 10^8^ ml^−1^ and 8.80 × 10^9^ ml^−1^ vs. 7.53 × 10^9^ ml^−1^) grew compared to predigested wheat, whereas *Fibrobacteres* was more abundant in latter cultures (1.32 × 10^7^ ml^−1^ vs. 2.87 × 10^7^ ml^−1^). Comparing predigested and native rye, significantly more *Firmicutes* and *Fusobacteria* (8.80 × 10^9^ ml^−1^ vs 7.29 × 10^9^ ml^−1^ and 1.46 × 10^8^ ml^−1^ vs. 0.305 × 10^8^ ml^−1^) grew on the predigested and significantly more *Proteobacteria* (1.19 × 10^9^ ml^−1^ vs. 2.38 × 10^9^ ml^−1^) grew on the native rye. Comparing predigested and native wheat, significantly more *Firmicutes* and *Spirochaetes* (7.53 × 10^9^ ml^−1^ vs. 6.27 × 10^9^ ml^−1^ and 3.02 × 10^8^ ml^−1^ vs. 1.61 × 10^8^ ml^−1^) grew on the predigested and significantly more *Proteobacteria* and *Actinobacteria* (1.02 × 10 ^9^ml^−1^ vs. 2.12 × 10 ^9^ml^− 1^ and 0.863 × 10^8^ ml^−1^ vs. 2.81 × 10^8^ ml^−1^) grew on native wheat.

**Figure 6.**
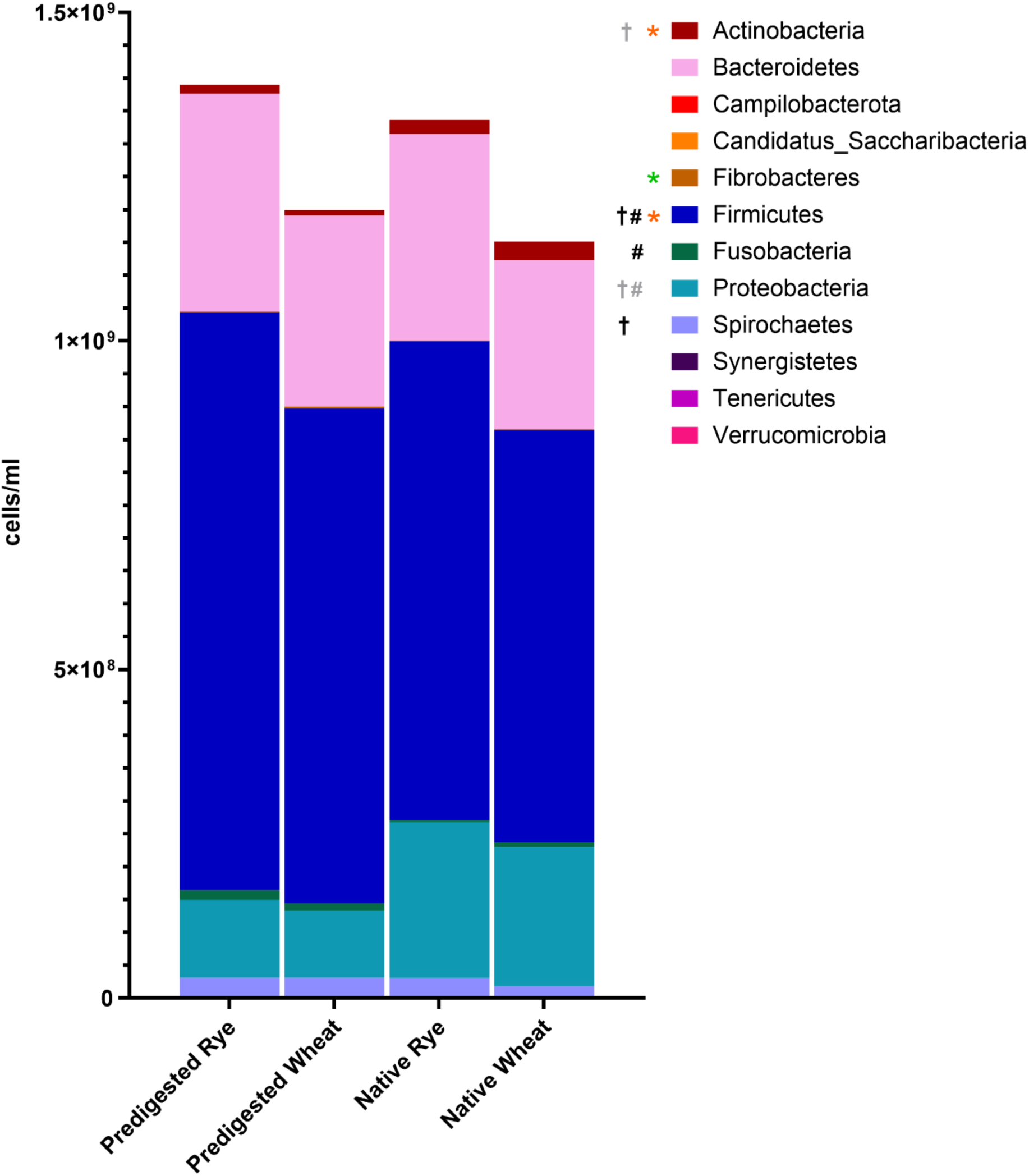
Absolute abundances of phyla after 24 hours of incubation of pig fecal microbiota (n=12) with predigested and native rye and wheat. * indicate significant differences between predigested rye and wheat (higher abundance for predigested rye with orange and higher abundance of predigested wheat with green). # indicate significant differences between predigested and native rye (higher abundance for predigested rye with black and higher abundance of native rye with grey). † indicate significant differences between predigested and native wheat (higher abundance for predigested wheat with black and higher abundance of native wheat with grey).

On the family level, many significant differences were found in the absolute abundances between the substrates (Figure 7). Looking into each substrate, for predigested rye the highest abundances were displayed by *Lachnospiraceae* (3.63 × 10^9^ ml^−1^), *Prevotellaceae* (2.81 × 10^9^ ml^−1^), *Streptococcaceae* (1.72 × 10^9^ ml^−1^) and *Ruminococcaceae* (1.15 × 10^9^ ml^−1^). For predigested wheat, the highest concentrations with were displayed by *Lachnospiraceae* (2.86 × 10^9^ ml^−1^), *Prevotellaceae* (2.38 × 10^9^ ml^−1^), *Clostridiaceae_1* (1.38 × 10^9^ ml^−1^) and *Ruminococcaceae* (1.29 × 10^9^ ml^−1^). For native substrates the highest abundances were found for *Prevotellaceae* (2.70 × 10^9^ ml^−1^ for rye and 2.03 × 10^9^ ml^−1^ for wheat), *Lachnospriaceae* (2.43 × 10^9^ ml^−1^ for rye and 1.56 × 10^9^ ml^−1^ for wheat), *Streptococcaceae* (1.45 × 10^9^ ml^−1^ for rye and 0.915 × 10^9^ ml^−1^ for wheat), *Succinivibrionaceae* (1.26 × 10^9^ ml^−1^ for rye and 0.964 × 10^9^ ml^−1^ for wheat) and *Clostridiaceae_1* (1.18 × 10^9^ ml^−1^ for rye and 1.80 × 10^9^ ml^−1^ for wheat).

**Figure 7.**
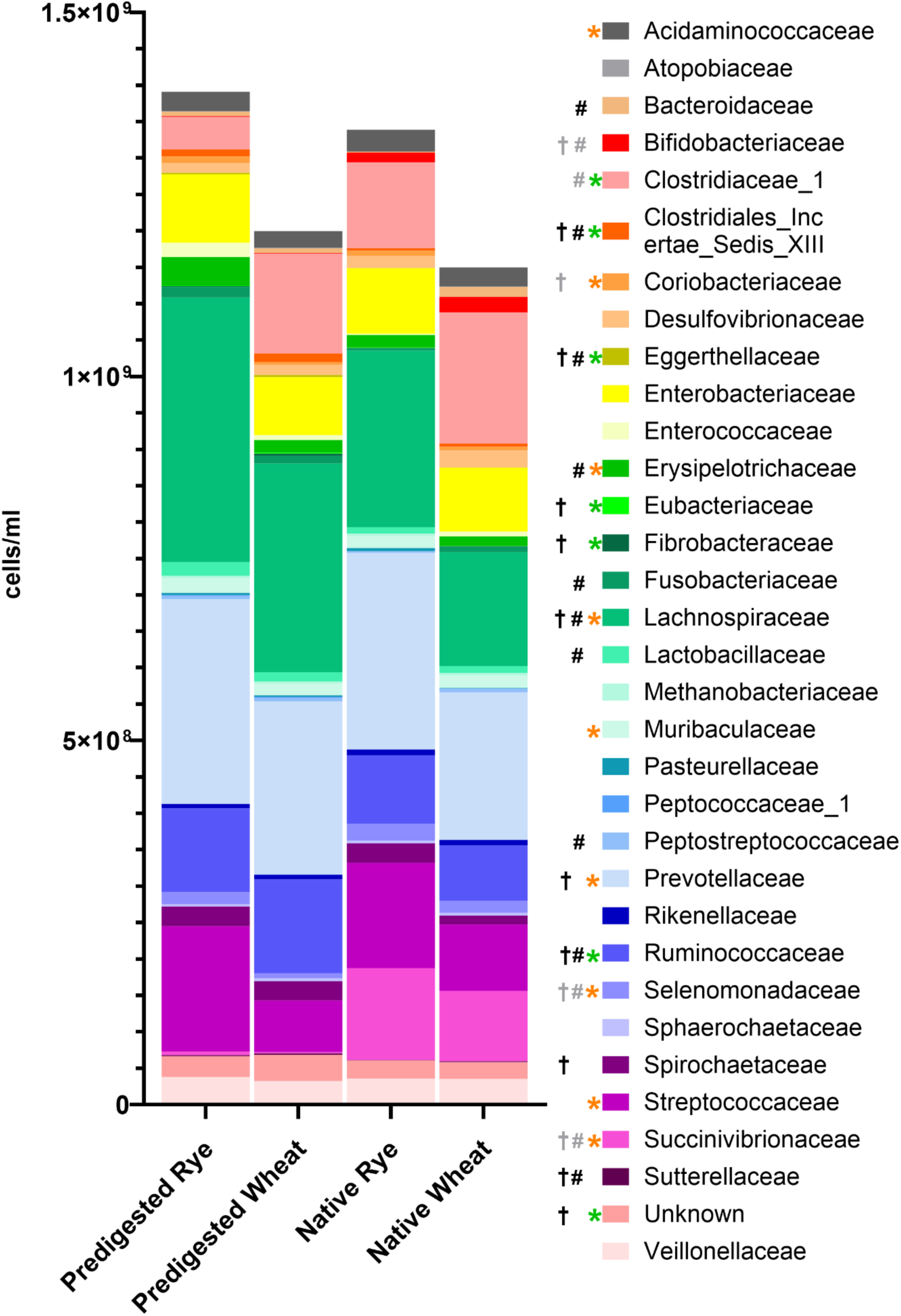
Absolute abundances of families after 24 hours of incubation of pig fecal microbiota (n=12) with predigested and native rye and wheat. * indicate significant differences between predigested rye and wheat (higher abundance for predigested rye with orange and higher abundance of predigested wheat with green). # indicate significant differences between predigested and native rye (higher abundance for predigested rye with black and higher abundance of native rye with grey). † indicate significant differences between predigested and native wheat (higher abundance for predigested wheat with black and higher abundance of native wheat with grey).

**Figure 8.**
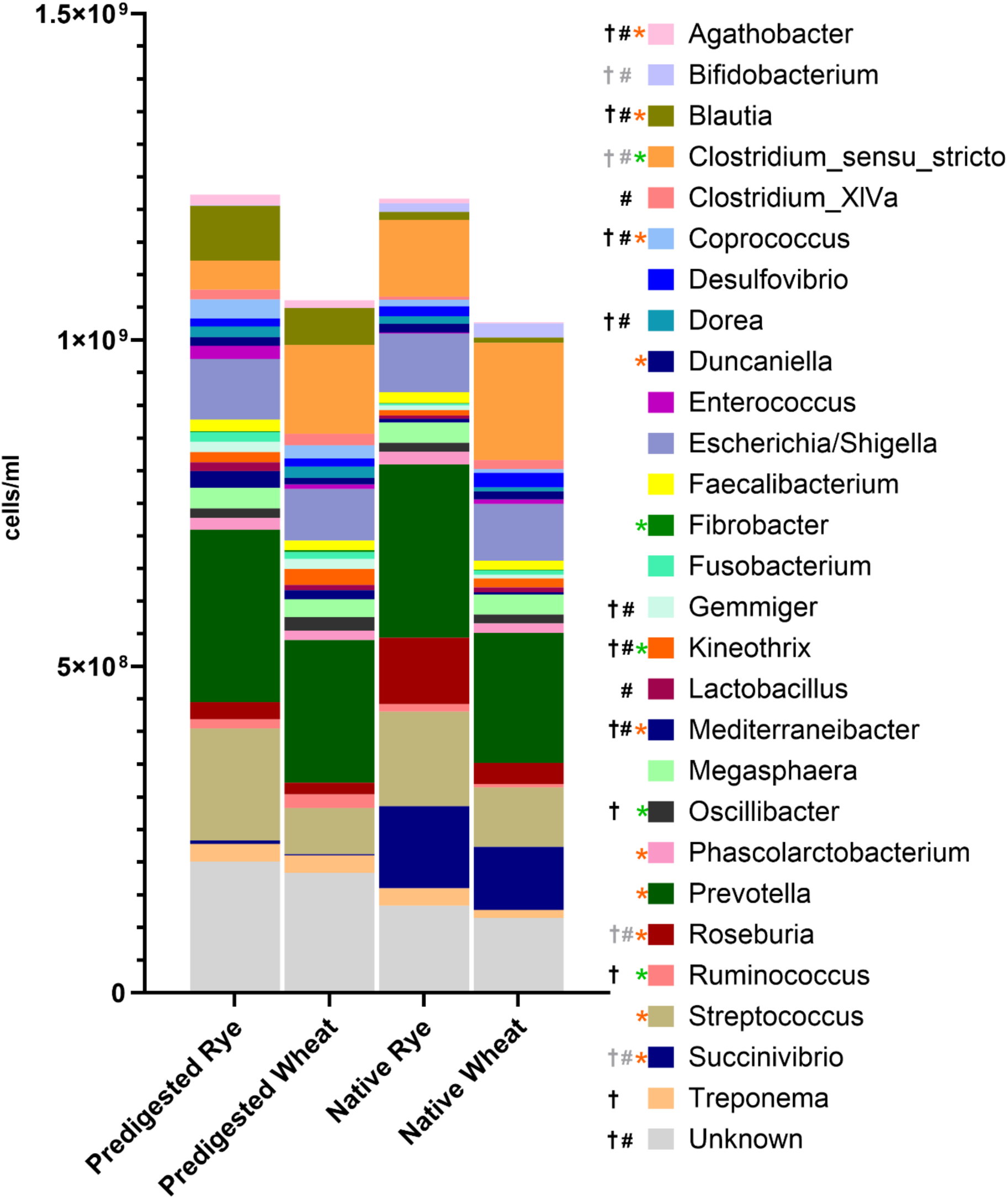
Absolute abundances of genera after 24 hours of incubation of pig fecal microbiota (n=12) with predigested and native rye and wheat. * indicate significant differences between predigested rye and wheat (higher abundance for predigested rye with orange and higher abundance of predigested wheat with green). # indicate significant differences between predigested and native rye (higher abundance for predigested rye with black and higher abundance of native rye with grey). † indicate significant differences between predigested and native wheat (higher abundance for predigested wheat with black and higher abundance of native wheat with grey).

On the genus level, many significant differences were found in absolute abundances between substrates (Figure 7, not all depicted). The highest abundances with predigested rye as a substrate were found for *Prevotella* (2.64 × 10^9^ ml^−1^), unknown genera (2.01 × 10^9^ ml^−1^) and *Streptococcus* (1.72 × 10^9^ ml^−1^). With predigested wheat as a substrate the highest abundances were found for *Prevotella* (2.19 × 10^9^ ml^−1^), unknown genera (1.84 × 10^9^ ml^−1^) and *Clostridium_sensu_stricto* (1.36 × 10^9^ ml^−1^). With native rye as a substrate the highest abundances were found for *Prevotella* (2.66 × 10^9^ ml^−1^), *Streptococcus* (1.45 × 10^9^ ml^−1^), unknown genera (1.33 × 10^9^ ml^−1^), *Succinivibrio* (1.25 × 10^9^ ml^−1^), *Clostridium_sensu_stricto* (1.17 × 10^9^ ml^−1^) and *Roseburia* (1.02 × 10^9^ ml^−1^). With native wheat as a substrate the highest abundances were found for *Prevotella* (1.99 × 10^9^ ml^−1^*), Clostridium_sensu_stricto* (1.79 × 10^9^ ml^−1^) and unknown genera (1.14 × 10^9^ ml^−1^).

Other genera with significant differences in abundance, that are not depicted in Figure 7, can be found in supplementary Table S1.

## Discussion

The aim of the present study was to investigate the composition of pig intestinal microbiota as well as their SCFA formation when grown on rye and wheat, respectively. Additionally, we evaluated the effects of predigestion of substrates through ileal cannulated minipigs on *in vitro* incubations.

Even though rye and wheat are closely related and display similar nutrient profiles, differences can be found in the content of amino acids, carbohydrates, minerals, enzyme activities and energy and also the digestibility of various components [9, 12, 25]. Differences in the nutrient profile of native cereals may therefore become even more pronounced during ileal digestion through different degradability of individual compounds, which was substantiated by our analysis in this study comparing native and predigested cereals. Those differences resulted in clear substrate-specific effects on all measured parameters in our *in vitro* experiments. The production of SCFA differed between predigested rye and wheat as well as between native and predigested substrates. More acetate was produced in cultures with predigested substrates and more from rye than from wheat. This indicates a higher energy yield for SCFA production in rye, especially for the predigested rye, which is most likely caused by the higher levels of fructan and soluble arabinoxylan levels in predigested rye than any of the other samples. Inulin and arabinoxylan have previously been found to promote propionate production [43, 44], which was also true for our experiments, where the highest amount of propionate was produced in cultures containing rye. Propionate can be used in the liver for gluconeogenesis [45] and has been found to induce the expression of satiety related hormones [46]. The SCFA has also been found to promote intestinal development and formation of jejunal tight junctions [22]. A propionate rich fermentation pattern can therefore be especially beneficial for pregnant sows that are fed restrictively [47] or weaned piglets, where weaning stress can disrupt intestinal barrier function [48]. Previous findings also indicated protective effects of propionate against *Salmonella* [49]. It has been shown before, that feeding rye leads to a higher amount of carbohydrates reaching the large intestine compared to wheat [9] along with reduced *Salmonella* shedding in an experimental infection [10] which might stem from differences in fermentation patterns as identified in our study. Contrary to a previous *in vivo* study, where rye and wheat diets were fed to pigs [50], the butyrate production was not found to be higher with rye compared to wheat in our study. Butyrate is important for colon homeostasis and function [51] and has also been shown to reduce *Salmonella* counts [52], therefore a butyrate rich fermentation pattern is regarded as beneficial for pigs. Especially arabinoxylans have previously shown a butyrogenic effect ins pigs *in vivo* [53] and we would therefore have expected a higher butyrate production on rye in our study as well. As *in vitro* experiments cannot depict all physiological processes like absorption from the intestine, the reason for the differences between *in vivo* studies and our experiments might lay in that complexity and cannot fully be explained here.

The butyrate production from native substrates was significantly higher than from the respective predigested substrates, which might be due to the overall higher substrate availability in native than predigested samples. Those results do, however, not reflect the substrate composition reaching the hindgut and again highlights the importance of predigestion to get realistic insights.

Bacterial concentrations and community composition after the growth on the different substrates demonstrated a clear separation between the native and the predigested forms, making it clear that results from native substrates in *in vitro* research cannot realistically depict *in vivo* microbial communities. Not only was the number of cells grown higher on the respective predigested substrates, but the community composition also differed notably. Comparing only communities grown on native substrates, a clustering based on individual pigs was observed. Growth on predigested substrates led to a notable differentiation between communities derived from rye and wheat, respectively. Looking more in detail into individual taxa many significant differences underline those findings. In the comparison of all four substrates, the highest abundance of the phylum *Firmicutes* was found when predigested rye was used as a substrate. Members of this phylum, especially from the families *Lachnospiracaeae* and *Ruminococcaceae,* are known to be butyrate and propionate producers [54]. *Lachnospiracaeae* were found to display the highest abundance of all four substrates in batches with predigested rye as a substrate, whereas *Ruminococcaceae* were found to display the highest abundance in batches containing predigested wheat. The native substrates yielded higher abundances of *Proteobacteria*, especially from the family *Succinivibrionaceae,* than predigested substrates. *Proteobacteria* were previously found to be higher abundant under an *E.* coli infection challenge in weaned piglets [55] and *Succinivibrionaceae* were associated with dysbiosis [56]. This indicates an also somewhat dysbiotic setting in the batches with native substrates. *Bifidobacteriaceae* were also higher abundant in cultures with native substrates, which may be due to the bifidogenic effect of high starch levels [57, 58]. High abundances of *Bifidobacteriaceae* were also found in settings with unnatural high substrate availability in the gut, for example in the condition of small intestinal bacterial overgrowth under untreated exocrine pancreatic insufficiency in a porcine model [59], with *Bifidobacteriaceae* from pig origin growing on a higher variability of substrates than those of human origin [58].

Known propionate producing bacteria that grew more on predigested substrates and more on rye than wheat are *Blautia, Mediterraneibacter* and *Phascolarctobacterium* [27, 37]. Those might be responsible for the higher propionate production in cultures with rye. In the case of butyrate, *Roseburia* and *Clostridium_sensu_stricto* were found to a higher extent when native substrates were used probably explaining the higher butyrate production in batches with native substrates [60]. Overall, predigested rye as a substrate created an environment for higher SCFA production and cell growth compared to predigested wheat, underlining the high value of rye as a component in pig diets to support health and well-being.

## Conclusion

Based on our results we conclude that, *in vitro,* rye as a substrate resulted in the production of higher amounts of the beneficial SCFA propionate compared to wheat which was accompanied by distinct community compositions between the two substrates. The study supports the view that rye can be used as a health promoting feed source in pig husbandry. We further conclude that in order to achieve *in vitro* results mimicking *in vivo* conditions as close as possible, a predigestion of substrates is crucial. Based on all parameters measured, native and predigested cereals yielded different results making the fistulated animal model essential for investigations described here. This aspect is not restricted to animal-based research but also applies for studies looking into *in vitro* substrate degradation using human material.

## Ethics approval

The animal trials were conducted in accordance with German regulations and approved by the Ethics Committee of Lower Saxony for the Care and Use of Laboratory Animals (LAVES: Niedersächsisches Landesamt für Verbraucherschutz und Lebensmittelsicherheit; reference: 33.8-42502-04-18/2884).

## Data Availability Statement

The datasets used and analysed during the current study are available from the corresponding author on reasonable request. All sequences are publicly available at the European Nucleotide Archive (PRJ coming soon).

## Authors’ contributions

A.v.F., C.B.H, C.V. and M.V. conceived the study. C.B.H. performed the experiments. C.V. provided the animal model. C.B.H., K.E.B.K. and S.W. performed laboratory analyses. M.V. performed bioinformatical analyses. C.B.H. and M.V performed data analysis and interpreted the results. C.B.H and M.V. wrote the manuscript. All authors reviewed and edited the manuscript and approved the final manuscript.

---

**Table S1.**
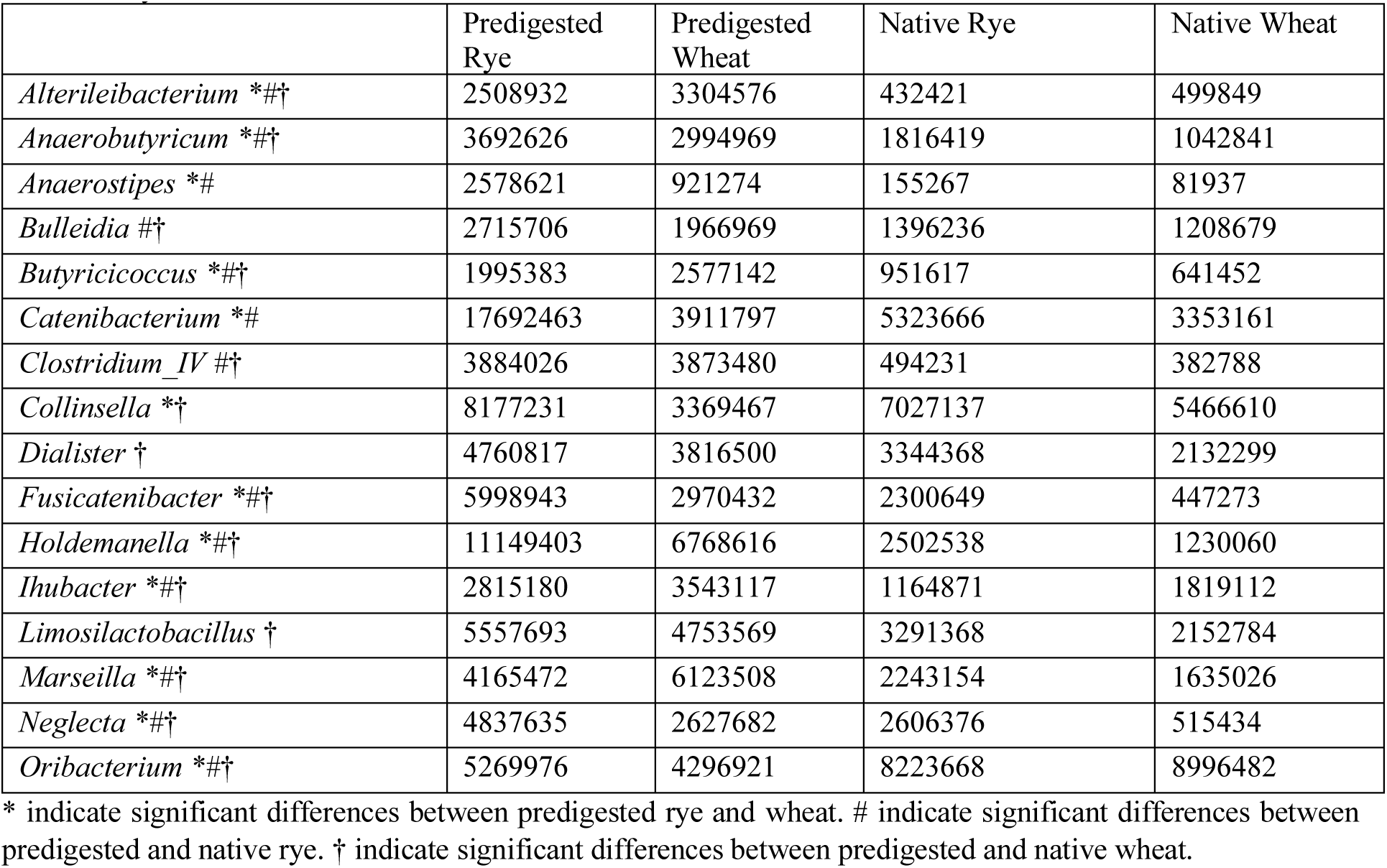
Genera with differences in absolute abundance (cells/ml) between growth on predigested und native wheat and rye.

|  | Predigested Rye | Predigested Wheat | Native Rye | Native Wheat |
| --- | --- | --- | --- | --- |
| *Alterileibacterium* *#† | 2508932 | 3304576 | 432421 | 499849 |
| *Anaerobutyricum* *#† | 3692626 | 2994969 | 1816419 | 1042841 |
| *Anaerostipes* *# | 2578621 | 921274 | 155267 | 81937 |
| *Bulleidia* #† | 2715706 | 1966969 | 1396236 | 1208679 |
| *Butyricicoccus* *#† | 1995383 | 2577142 | 951617 | 641452 |
| *Catenibacterium* *# | 17692463 | 3911797 | 5323666 | 3353161 |
| *Clostridium_IV* #† | 3884026 | 3873480 | 494231 | 382788 |
| *Collinsella* *† | 8177231 | 3369467 | 7027137 | 5466610 |
| *Dialister* † | 4760817 | 3816500 | 3344368 | 2132299 |
| *Fusicatenibacter* *#† | 5998943 | 2970432 | 2300649 | 447273 |
| *Holdemanella* *#† | 11149403 | 6768616 | 2502538 | 1230060 |
| *Ihubacter* *#† | 2815180 | 3543117 | 1164871 | 1819112 |
| *Limosilactobacillus* † | 5557693 | 4753569 | 3291368 | 2152784 |
| *Marseilla* *#† | 4165472 | 6123508 | 2243154 | 1635026 |
| *Neglecta* *#† | 4837635 | 2627682 | 2606376 | 515434 |
| *Oribacterium* *#† | 5269976 | 4296921 | 8223668 | 8996482 |

\* indicate significant differences between predigested rye and wheat. # indicate significant differences between predigested and native rye. † indicate significant differences between predigested and native wheat.

